## Supplement Figures for "Single-cell characterization of human GBM reveals regional differences in tumor-infiltrating leukocyte activation"

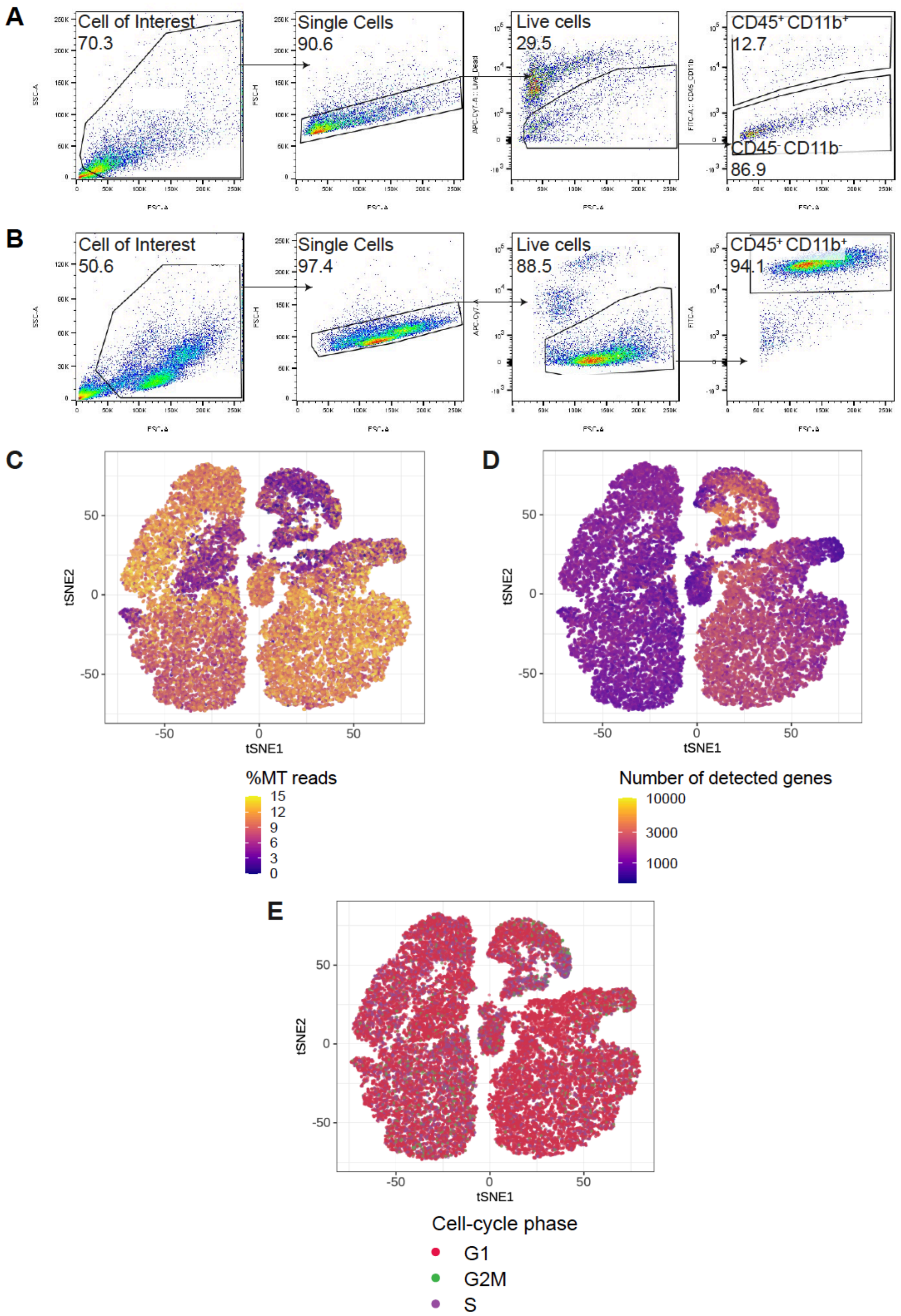


**Figure 1 – figure supplement 1. CD45+ CD11b+ immune cells gating strategy and quality control of scRNA-seq data.**

**(A, B)** Gating strategy for paired tumor-derived (**A**) and PBMCs (**B**); after debris, doublet and dead cell removal, immune cells were assessed as CD45+ and/or CD11b+. **(C-E)** Percentage of mitochondrial (MT) reads (**C**) number of detected genes (**D**) and cell-cycle phase (**E**) overlaid on tSNE representation. Please see Supplementary File 2 for quantification of **C-E**.

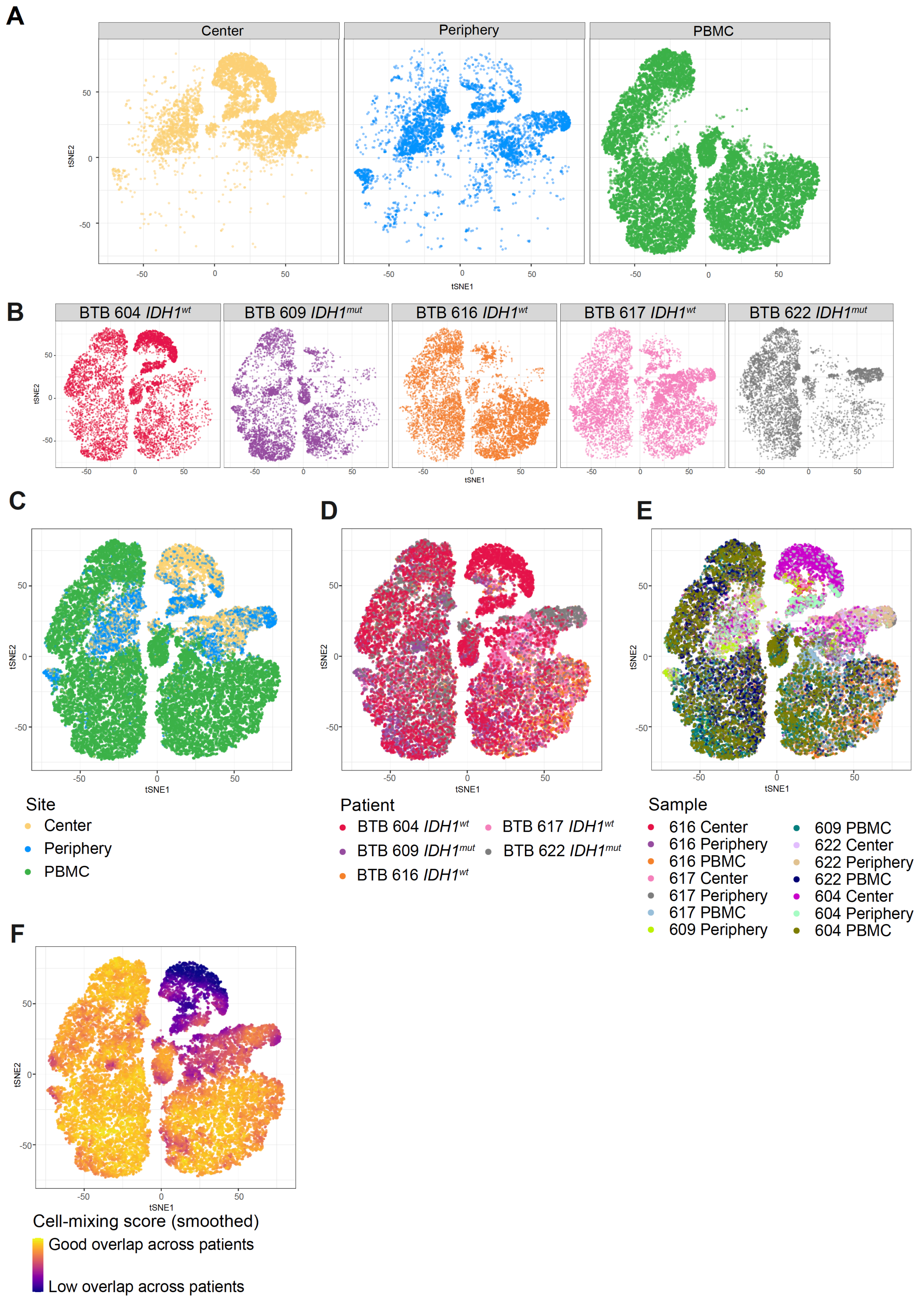


**Figure 1 – figure supplement 2. Patient representation among clusters.**

**(A, B)** tSNE map stratified according to site (**A**) and patient (**B**). (**C-E**) tSNE maps showing all cells, colored by site (**C**), patient (**D**) and sample (**E**). (**F**) Cell-mixing score [69] overlayed on tSNE representation.

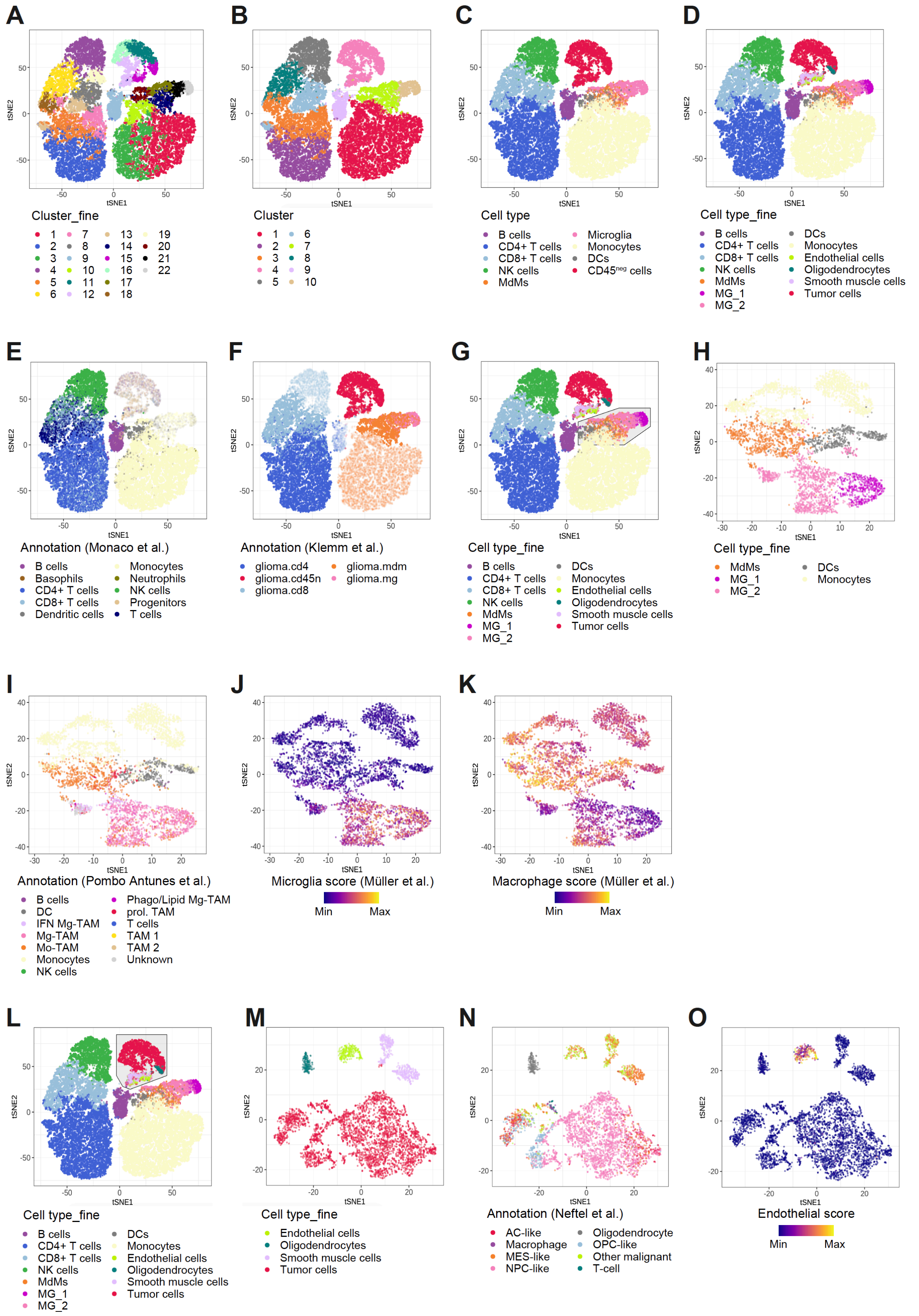


1100

1101 **Figure 2 – figure supplement 1. Cross-referencing scRNA-seq data with published da-**

1102 **tasets.**

**(A-D)** Using hierarchical clustering, identified cell clusters at different levels of granularity (**A, B**) which were then annotated into nine distinct cell types for the immune subset and four cell types for the CD45neg subset (**C, D**). (**E, F)** Immune cell types were annotated by referencing to a dataset of bulk RNA-seq samples of sorted immune cell types from human PBMC (**E**) [16] and MdMs and microglia were annotated by comparing to a dataset of bulk RNA-seq samples of sorted immune cell types from the tumor microenvironment of human gliomas (**F**) [5]. Clusters are highlighted which were annotated using each respective reference dataset. (**G-K**) Subset analysis on tumor innate immune cells (**G**), showing re-clustering on the subset (**H**), annotation by referencing to a 10X genomics scRNA-seq dataset of TAMs from the GBM TME of 7 newly diagnosed human patients (**I**) [9] and signature scores defined from scRNA-seq of glioma TAMs (**J, K**) [7]. (**L-O**) Subset analysis on CD45$^{neg}$ cells (**L**), showing re-clustering on the subset (**M**), annotation by whole-transcriptome comparison to a scRNA-seq dataset of *IDH1*$^{wt}$ GBM (**N**) [17] and an endothelial score which was defined by averaging the center and scaled expression levels of the genes CDH5, VWF, CD34 and PECAM1 (**O**).

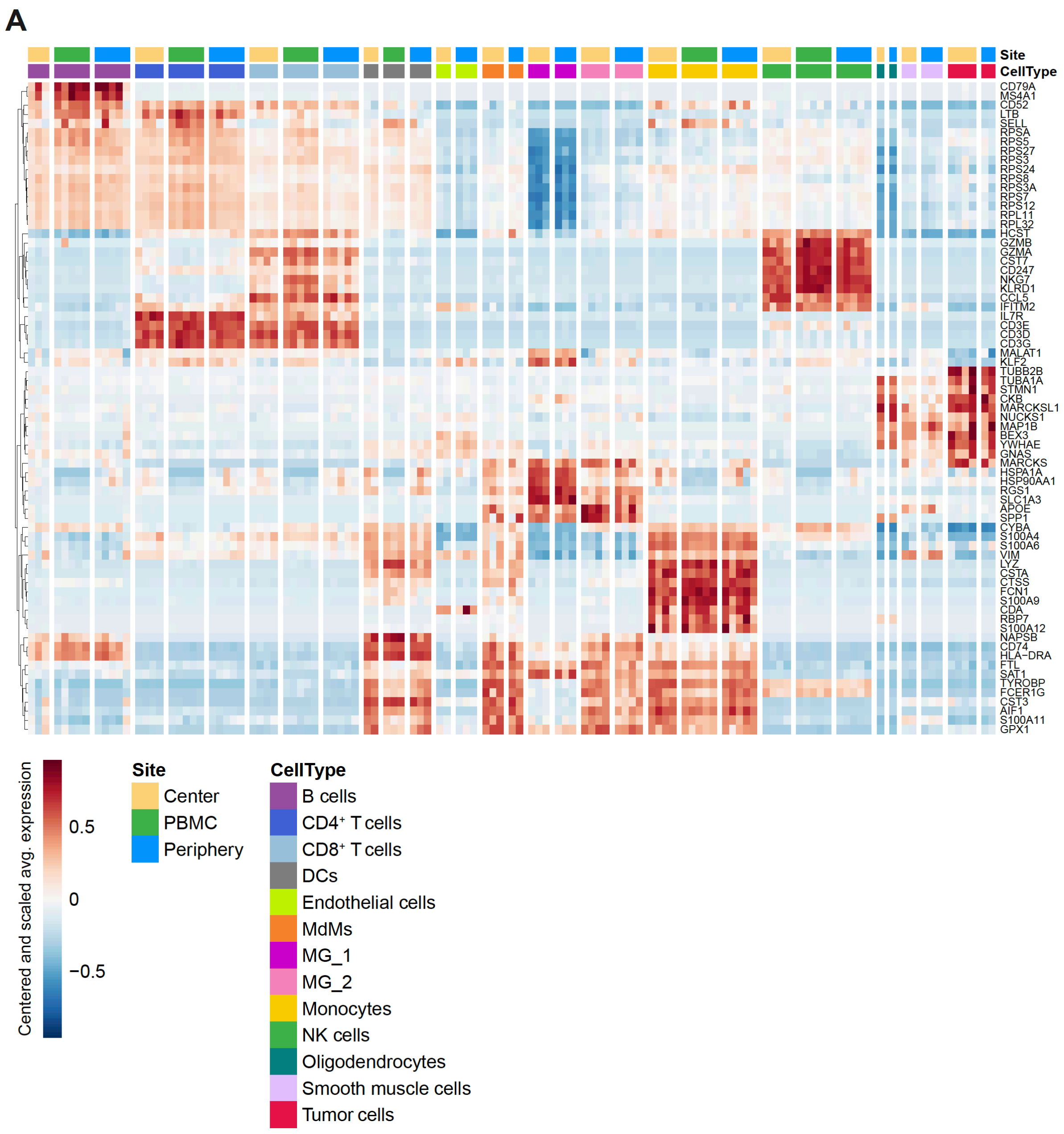


**Figure 2 – figure supplement 2. Cell type specific gene expression.**

**A,** Heatmap displaying genes whose expression is most specific to each cell type. Columns are ordered by site and cell type, and rows show centered and scaled expression values, hierarchically clustered.

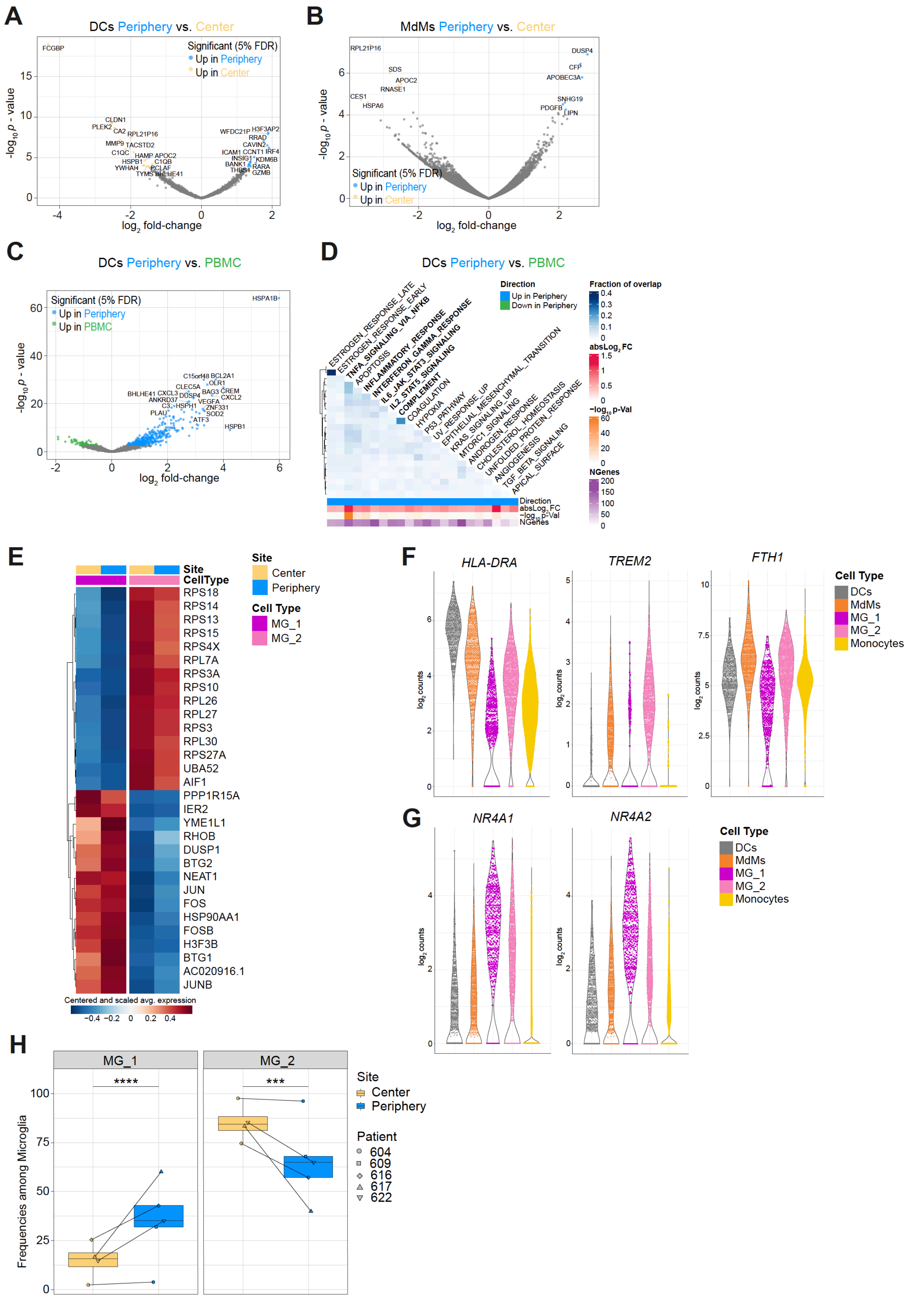


**Figure 3 -figure supplement 1. Regionally dependent transcriptional profiles of innate immune subsets and MG subclusters MG_1 and MG_2.**

**(A, B)** Volcano plots showing differentially expressed genes (FDR corrected *p* value < 0.05, indicated by colors) in DCs **(A)** and MdMs **(B)** from tumor periphery versus center. **(C)** Volcano plot showing differentially expressed genes (FDR corrected *p* value < 0.05, indicated by colors) in DCs from tumor periphery versus PBMC. **(D)** Heatmap representation of GSEA between DCs from tumor periphery and PBMC using Hallmark collection of major biological categories. **(E)** Heatmap displaying the cluster-specific genes identifying MG_1 and MG_2 subclusters. Columns are ordered by site and cell type, and rows show centered and scaled normalized average expression values, hierarchically clustered. **(F, G)** Violin plots showing average expression levels ($\log_2$ counts) of selected markers among mononuclear phagocyte populations. **(H)** Frequencies of MG_1 and MG_2 subpopulations among total microglia between center and periphery. Symbols represent individual patients and paired samples are indicated by connecting lines. Statistical significance was assessed by diffcyt-DA-voom method, ***FDR corrected p value $p \leq 0.001$, ****FDR corrected p value $\leq 0.0001$.

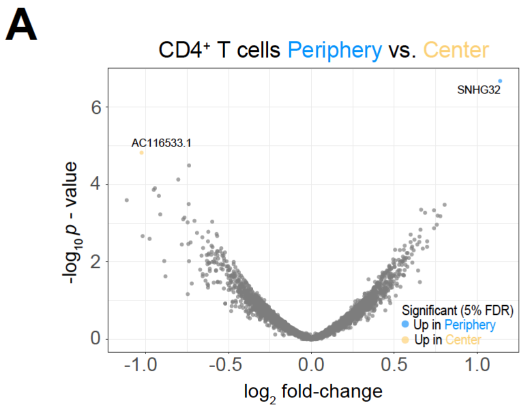


**Figure 4 – figure supplement 1. Differential expression analysis between tumor center and peripheral CD4+ T cells.**

(**A**) Volcano plot showing differentially expressed genes (FDR corrected p value < 0.05, indicated by blue and yellow) in CD4+ T cells from tumor periphery versus tumor center.

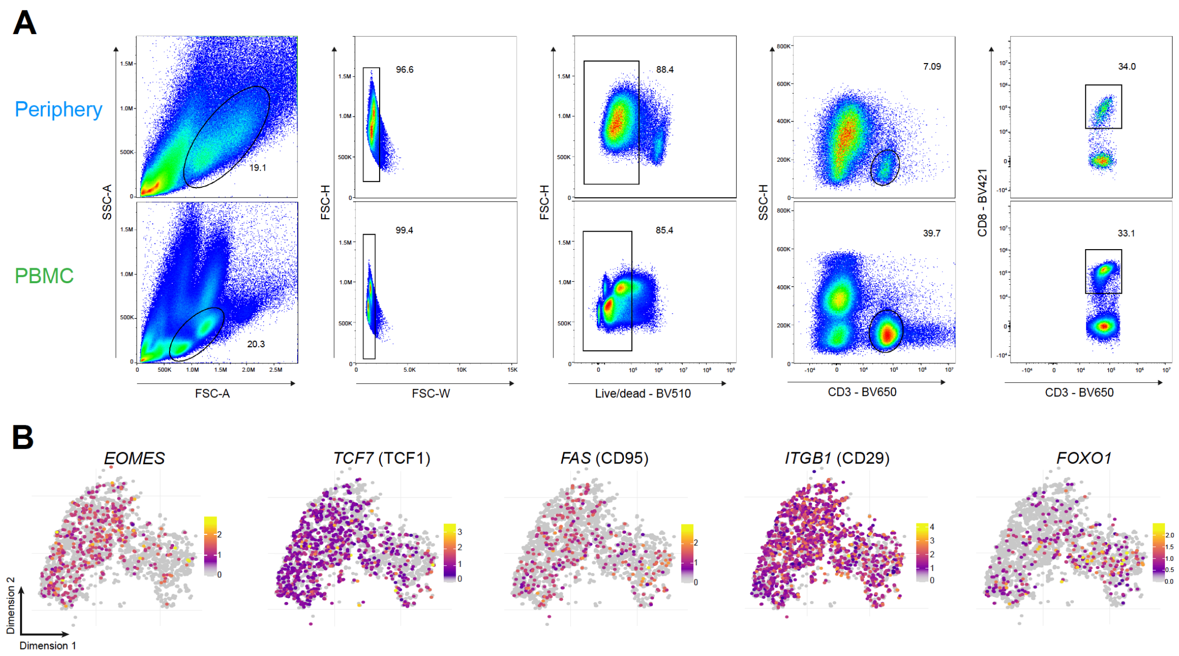


**Figure 5 – figure supplement 1. Phenotypic characterization of PBMC CD8$^{+}$ T cells.** (**A**) Gating strategy for paired tumor-periphery and PBMC cells; after debris, doublet and dead cell removal, CD8+ T cells were identified as CD3+ CD8+ events. (**B**) Single-cell expression of additional markers associated with naïve/memory.

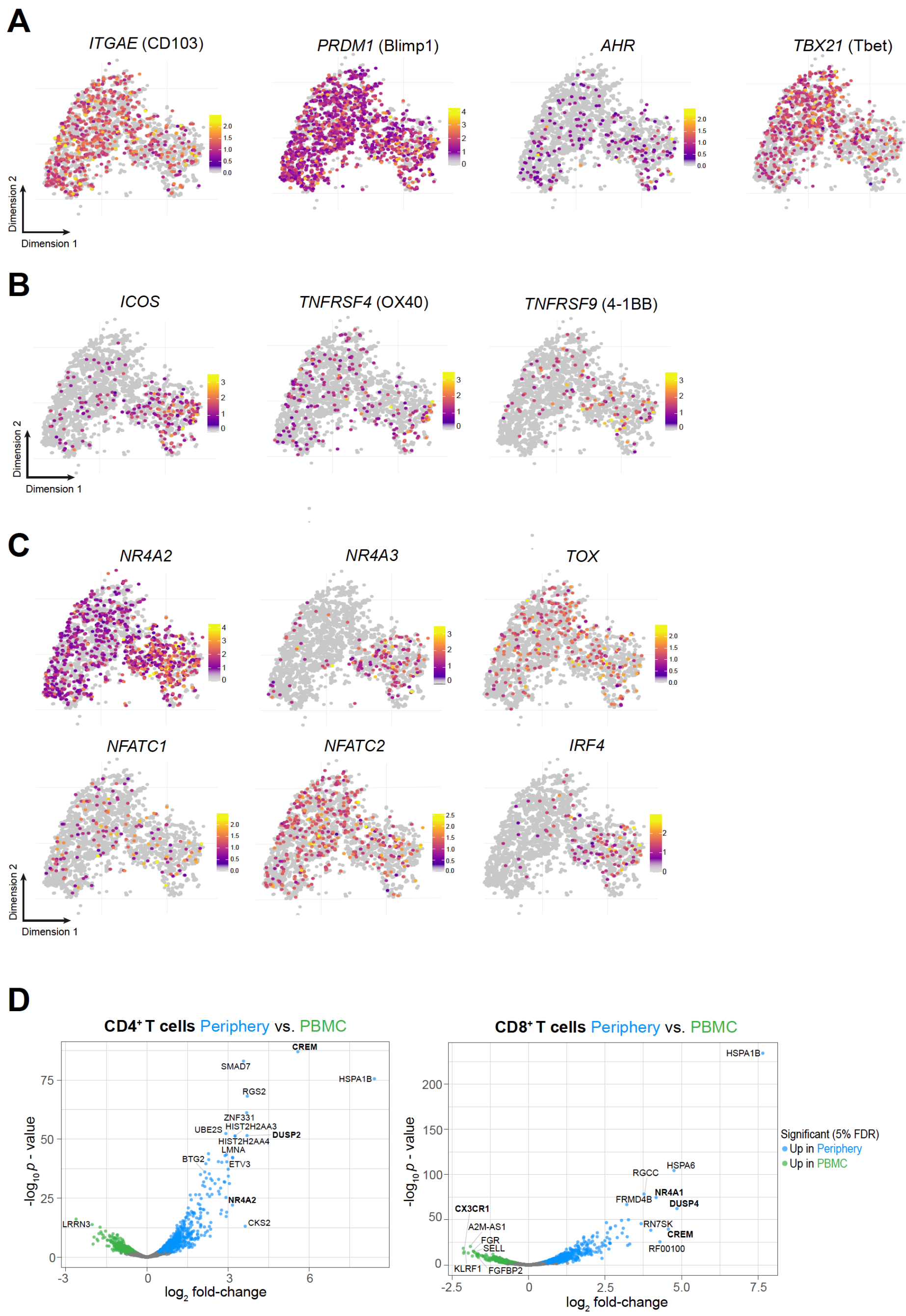


**Figure 6 – figure supplement 1. Phenotypic characterization of Periphery CD8$^+$ T cells.** (**A-C**) Single-cell expression of additional markers associated with tissue-resident memory (A), T cell co-stimulation (B) and T cell exhaustion/dysfunction (C) overlaid on tSNE CD8+ T cell cluster. (**D**) Volcano plot showing differentially expressed genes (FDR corrected p value < 0.05, indicated by blue and green) in CD4$^+$ (left) and CD8$^+$ T cells from tumor-periphery versus PBMC.

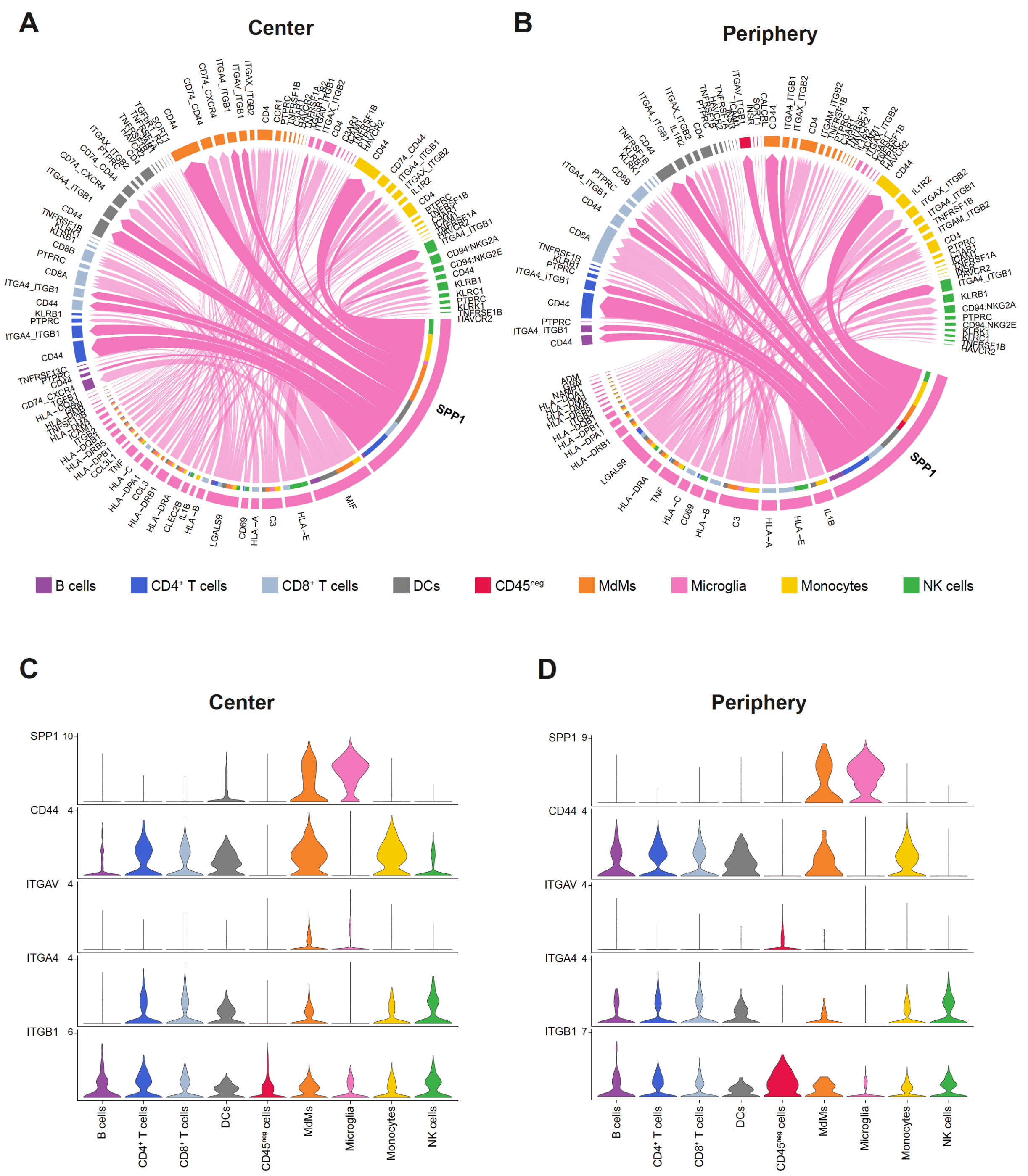


**Figure 7 – figure supplement 1. Cell-cell communication analysis using CellChat.**
(**A, B**) Chord diagram showing significant interactions from microglia to all cell clusters in center (**A**) and periphery (**B**). The inner bar colors represent the targets that receive signal from the corresponding outer bar. The inner bar size is proportional to the signal strength received by the targets. Chords indicate ligand-receptor pairs mediating interaction between two cell clusters, size of chords is proportional to signal strength of the given ligand-receptor pair.
(**C, D**) Violin plots showing the expression distribution of signaling genes (in $\log_2$ counts) involved in the inferred SPP1 signaling network in center (**C**) and periphery (**D**).
